# LuxRep: a technical replicate-aware method for bisulfite sequencing data analysis

**DOI:** 10.1101/444711

**Authors:** Maia Malonzo, Viivi Halla-aho, Mikko Konki, Riikka J. Lund, Harri Lähdesmäki

## Abstract

DNA methylation is measured using bisulfite sequencing (BS-seq). Bisulfite conversion can have low efficiency and a DNA sample is then processed multiple times generating DNA libraries with different bisulfite conversion rates. Libraries with low conversion rates are excluded from analysis resulting in reduced coverage and increased costs. We present a method and software, LuxRep, that accounts for technical replicates from different bisulfite-converted DNA libraries. We show that including replicates with low bisulfite conversion rates generates more accurate estimates of methylation levels and differentially methylated sites.

**Availability:** An implementation of the method is available at https://github.com/tare/LuxGLM/tree/master/LuxRep

## 1 Introduction

In the optimal case, the bisulfite conversion rate of a DNA library is high (above 99%). However, when an experiment yields a low conversion rate the common lab practice is to exclude the DNA library so as to avoid overestimation of methylation levels, resulting in additional costs or reduced data depending on whether a replacement library is prepared or not. An advanced computational approach to handle poor conversion rates would render exclusion of samples unnecessary. The methylation analysis method LuxGLM (Äijö *et al*., 2016) estimates methylation levels from bisulfite sequencing data using a probabilistic model that accounts for bisulfite conversion rate. Though the previous model was able to handle biological replicates with a general linear model component, it assumed data from each sample consisted of only a single bisulfite-converted DNA library. In this work we propose LuxRep, an improved method and software to allow use of replicates from different DNA libraries with varying bisulfite conversion rates. To make LuxRep tool computationally efficient and thus more applicable to genome-wide analysis we also propose to use variational inference.

## 2 Methods

We start by briefly reviewing the underlying statistical model (Äijö *et al*., 2016) and then introduce our extension that can handle technical replicates. Briefly, the conditional probability of a sequencing readout being “C” in BS-seq data is a function of the experimental parameters that include sequencing error (seq_err_) and bisulfite conversion rate (BS_eff_), and depends on the methylation level *θ* ∈ [0, 1]. If a read was generated from an unmethylated cytosine (C), the conditional probability *p*_BS_(“C”*|*C) is given by (1 *-* BS_eff_)(1 *-* seq_err_) + BS_eff_ seq_err_. The observed total number of “C” readouts for a single cytosine is binomially distributed, *N*BS,C ~ Bin(*N*BS*, p*BS(“C”)), where *N*_BS_ is the total number of reads and *p*_BS_(“C”) = *p*_BS_(“C”*|*5mC) *θ*+ *p*_BS_(“C”*|*C)(1–*θ*). Finally, LuxGLM uses a general linear model with matrix normal distribution to handle covariates; see (Äijö *et al*., 2016) for further details.

We extend the model to allow modelling of technical replicates wherein the methylation level *θ* is the same for all different bisulfite-converted DNA libraries from the same biological sample but the experimental parameters (seq_err_ and BS_eff_) vary. In the modified model (Figs. 1a and S1a), *N*_BS,C_ and *N*_BS_ represent the observed “C” and total counts, respectively, from each of the *M_i_* technical replicates per biological sample *i* ∈ {1*,.., N*}. Note that the experimental parameters BS_eff_ and seq_err_ are sample and replicate-specific. See Supplementary Materials for further details.

**Figure 1.**
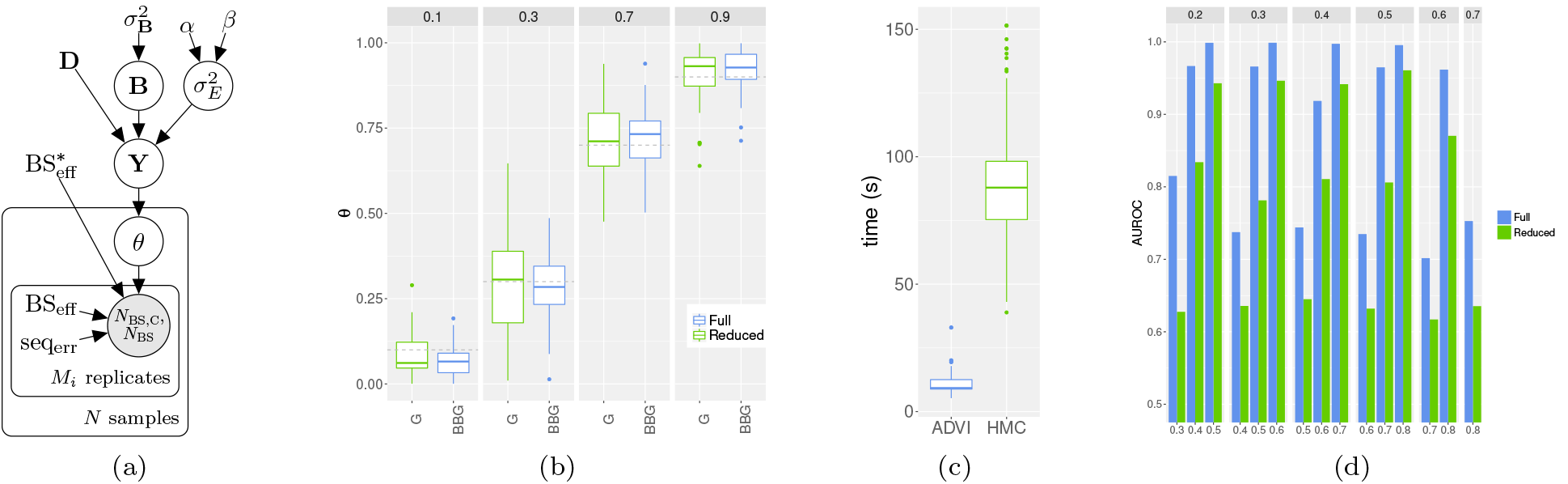
(a) Plate diagram of the LuxRep model for estimating methylation level of a single cytosine with technical replicates. The circles represent latent (white) and observed (gray) variables and the unbordered nodes represent constants. (b) Methylation levels estimated using the full and reduced LuxRep models with varying methylation level *θ* with input data from one good (“G”) and two bad (“B”) replicates (for the full model) and one “G” replicate (for the reduced model). Boxplots of posterior means from *n* = 100 independent simulated data sets are shown. Number of reads was 12 for each replicate and model was estimated using ADVI. (c) Comparison of running times using ADVI and HMC for model evaluation and including post-processing of output files. Input data used described in (b). (d) Accuracy in identifying differential methylation using the full and reduced LuxRep models measured by AUROCs based on 200 positive (Δ*θ≠* 0) and 200 negative (Δ*θ=* 0) datasets. Two groups (1 and 2) with four biological replicates and each biological replicate with technical replicates as described in (b) with varying methylation difference and levels (*θ*_1_ in top panel and *θ*_2_ in x-axis).

Äijö *et al*. (2016) used Hamiltonian Monte Carlo (HMC) for model inference, whereas in variational inference (VI) the posterior *p*(*ϕ|***X**) of a model is approximated with a simpler distribution *q*(*ϕ*;*ρ*),which is selected from a chosen family of distributions by minimizing divergence between *p*(*ϕ|***X**) and *q*(*ϕ*;*ρ*). We use the automatic differentiation variational inference algorithm (ADVI) from Kucukelbir *et al*. (2015), which is integrated into Stan. ADVI is used to generate samples from the approximative posterior *q*(*ϕ*;*ρ*).

Our software consists of two modules: 1) estimation of experimental parameters from control data, and 2) inference of methylation level and differential methylation using the fixed experimental parameters.

## 3 Results

We simulated technical replicates with low (BS_eff_= 0.9) and high (BS_eff_ = 0.995) BS conversion rates (BCRs) with varying sequencing depth *N*BS and methylation level (*θ* ∈ [0.1, 0.9]). Two scenarios were simulated consisting of three technical replicates each: (i) two replicates with high BCR (i.e. good samples, ‘G’) and one with low BCR (i.e. bad sample, ‘B’), and (ii) one ‘G’ replicate and two ‘B’ replicates. Each scenario was analyzed using (i) the *full* LuxRep model and (ii) a *reduced* model with experimental parameters fixed to BS_eff_ = 1, seq_err_ = 0 and 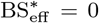, and using the “C” and “T” counts from only the ‘G’ samples to simulate the traditional approach of not accounting for experimental parameters. Results from estimating the models with HMC and ADVI were also compared.

The variance of the estimates using the full model was generally lower across *θ* and *N*_BS_ values (Figs. 1b and S2a) demonstrating the utility of using LuxRep with replicates of varying BCRs. The decrease in variance was generally greater with the second scenario. Improvements in the estimates were comparable when using HMC and ADVI (Fig.S2a), with significant reduction in running times with the latter (Fig. 1c).

To test the utility of LuxRep in determining differential methylation with replicates with low BCRs the experimental setups described above were extended to simulate two groups with four biological samples each (with each biological sample consisting of the technical replicates) and where the two groups had varying methylation difference Δ*θ* (0.1, 0.2 and 0.3). The full LuxRep model was more accurate at identifying differential methylation than the reduced model as shown by the higher area under the ROC curves (AUROCs) when using either ADVI (Fig. 1d) or HMC (Fig. S3). The improvement in AUROCs is notable with smaller differential methylation (i.e..6 Δ*θ*= 0.1). The increase in AUROC was also more pronounced with the second scenario.

To test the utility of LuxRep on an actual bisulfite sequencing dataset, methylation levels were estimated from an RRBS dataset consisting of two individuals and three replicates each (two low and one high BCR). The replicate with high BCR was analyzed with the full model while the two low-BCR replicates were analyzed with both the full and reduced models. The difference in the estimated methylation levels (1000 CpG sites) between the high-BCR replicate and the low-BCR replicates using the full and reduced models were measured by taking their euclidean distance which showed greater similarity when using the full model (individual 1: reduced - 2.55, full - 2.49; individual 2: reduced - 2.29, full - 2.23).

## 4 Conclusions

LuxRep tool described in this paper allows technical replicates with varying bisulfite conversion efficiency to be included in the analysis. LuxRep improves the accuracy of differential methylation analysis and lowers running time of model-based DNA methylation analysis.

## Acknowledgements and Funding

This work has been supported by the Academy of Finland Centre of Excellence in Molecular Systems Immunology and Physiology Research. We acknowledge the computational resources provided by the Aalto Science-IT project and the Finnish Functional Genomics Centre and Biocenter Finland is acknowledged for the infrastructure support.

## Supplementary information

### 1 LuxRep model

LuxRep is organised into two modules for estimating (1) experimental and (2) biological parameters from DNA bisulfite sequencing data. For estimation of biological parameters, LuxRep retains the general linear model component of LuxGLM (see Fig. 1a in main text) where the variables **Y**, **D**, **B** and **E** refer to, respectively, the unnormalized methylation fractions, design matrix, parameter matrix and noise. To emphasize the new features, the level for only one methylation modification (methylcytosine, 5mC) is included in this work.

The data consists of *N* biological samples (*i* ∈ {1*,…, N*}), each of which has *M_i_* technical replicates corresponding to different bisulfite-converted DNA library preparations. Samples prepared for BS-seq are typically spiked-in with unmethylated control DNA (often Lambda phage genome) that allows estimation of bisulfite conversion efficiency BS_eff_. The LuxGLM model (Äijö *et al*., 2016) was modified to determine experimental parameters for each technical replicate separately (shown as the “replicates” plate in the diagram in Fig. S1a). The circles represent latent (white) and observed (gray) variables and the squares/unbordered nodes represent fixed values (for parameters and hyperparameters).

**Figure S1:**
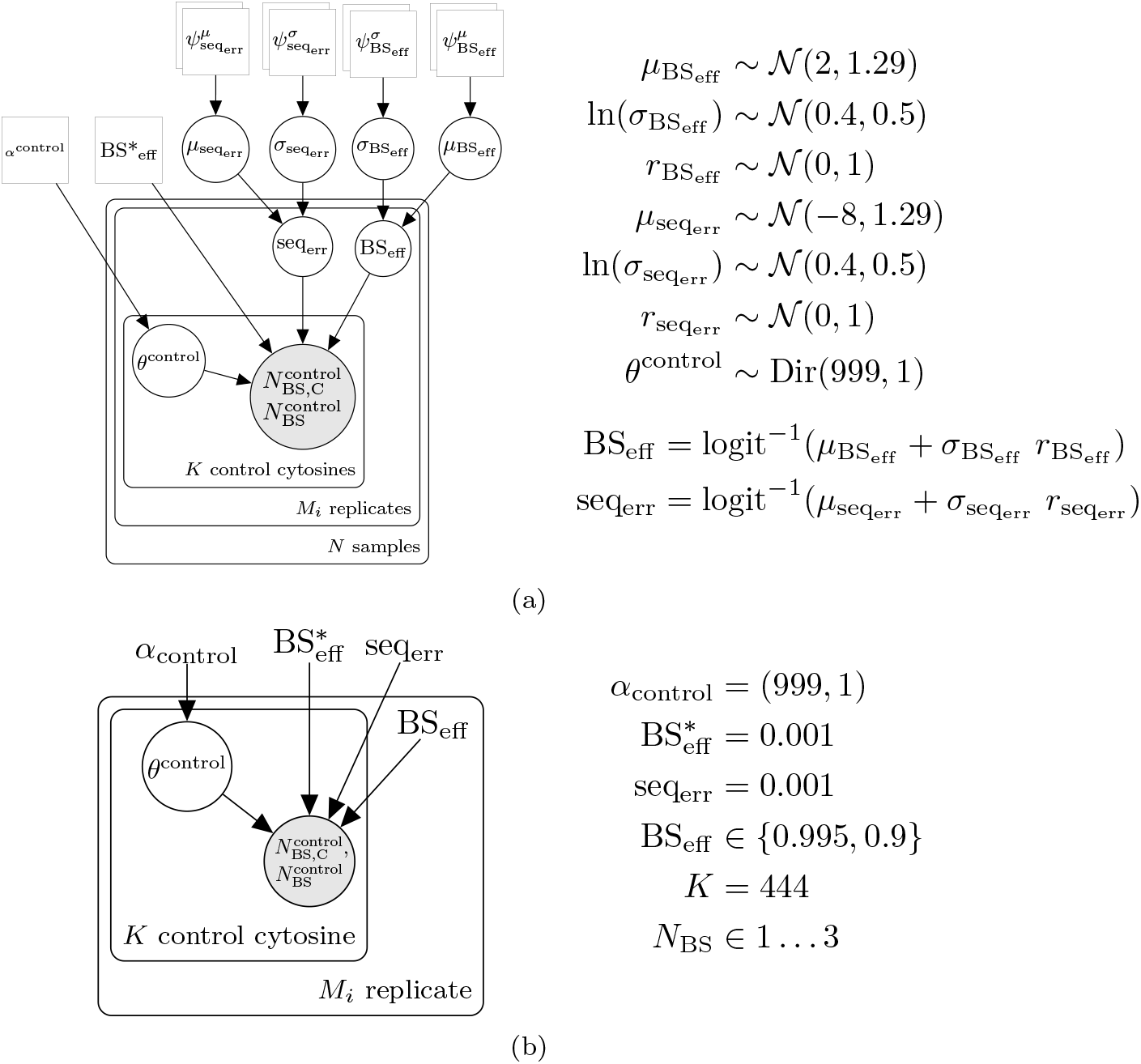
Plate diagrams of LuxRep model for the module analyzing experimental parameters from control data (a) and model generating dummy control data (b).

Incorrect bisulfite conversion rate, 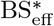, was set to a fixed value (0.1%) (in LuxGLM it was estimated from control data) because genome scale bisulphite sequencing typically do not include methylated cytosine control data.

To facilitate genome wide analysis, in our model implementation the experimental parameters are first computed from the control data since all cytosines per replicate have the same value for these parameters. Methylation levels are then determined individually for each cytosine, and differential methylation thereafter, using the pre-computed experimental parameters as fixed input.

### 2 Estimating experimental parameters

Dummy control cytosine data were generated using the model illustrated in Fig. S1b. Based on a cursory examination of an actual dataset generated from spiked-in Lambda phage DNA (data not shown), bisulfite sequencing data for 444 control cytosine were simulated with number of reads per cytosine *N*_BS_ ∈1 … 3. Experimental parameters were set to fixed values while the methylation modification fractions *θ*^control^ were drawn from Dir(z) (parameters listed in Fig. S1b).

Sequencing error and bisulfite conversion rates were estimated using the model illustrated in the plate diagram in Fig. S1a with priors and hyperpriors used listed beside.

### 3 Estimating methylation level and comparing running time

Datasets (*n* = 100) were analysed with the full and reduced LuxRep model with varying methylation levels (columns, values shown at topmost panel), varying number of reads (rows, values shown on right panel), different combinations of replicates with varying BCRs (‘G’ and ‘B’) (x-axis), and using either HMC or ADVI to evaluate or approximate the posterior, respectively S2b). The datasets were generated using the model illustrated in Fig. S2a with methylation levels and experimental parameters randomly generated with the beta distribution with parameters set to values listed to the right of the diagram. Simulations consisted of two subsets with differing combination of BCRs, ‘GGB’ and ‘GBB’.

**Figure S2:**
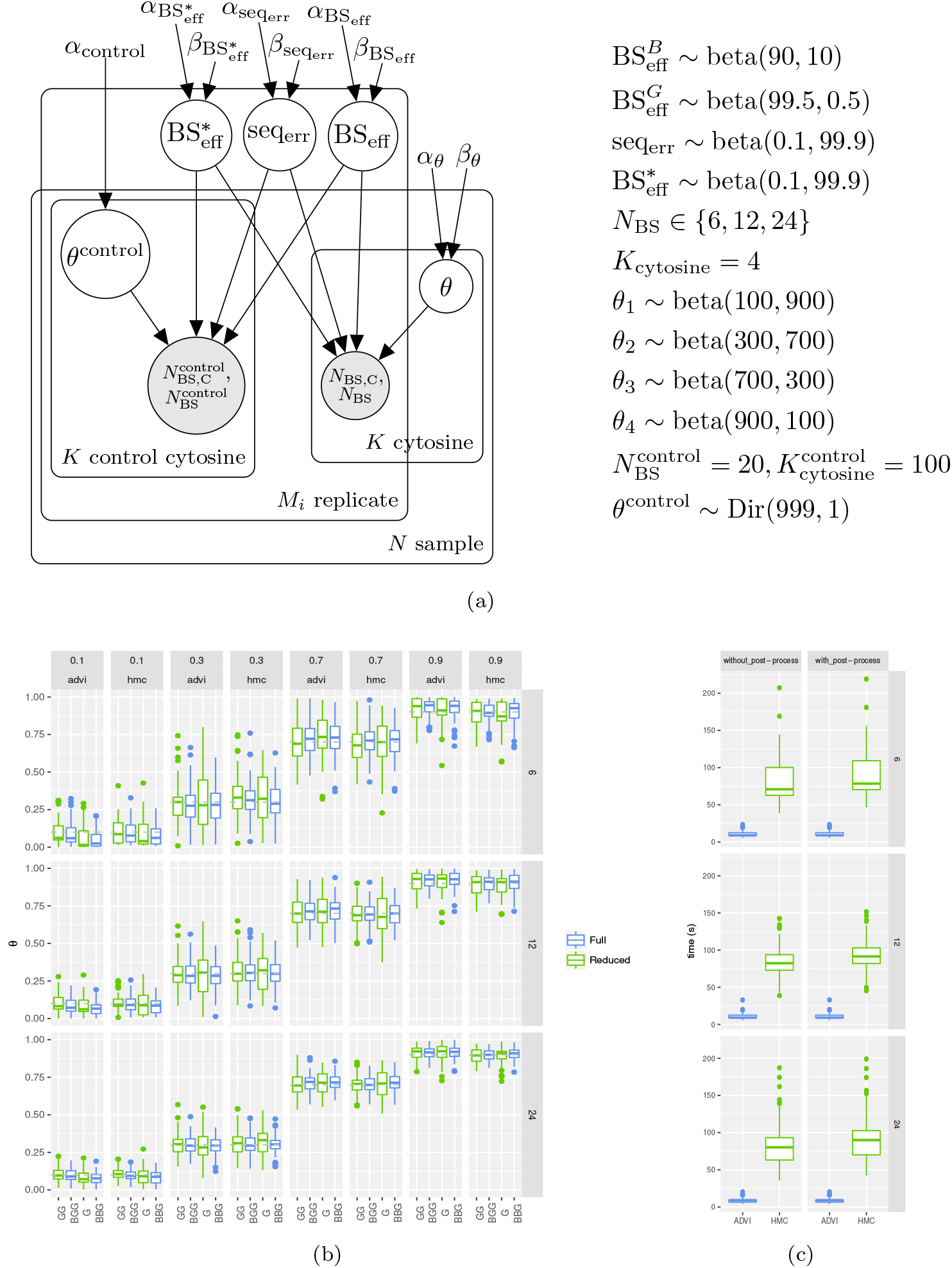
(a) Plate diagram of model for generating dummy data for estimating methylation level, (b) boxplots of estimates of methylation levels, and (c) comparison of running times using HMC and ADVI for model evaluation.

For each simulated data set we estimated the methylation level *θ* using the posterior mean of samples (*S* = 1000) drawn from the posterior (HMC) and approximate (ADVI) posterior distribution. Fig. S2b shows boxplots of the posterior means (*n* = 100). The hyperpriors used are listed below Fig. S2a.

Running times were measured using the Stan (Carpenter *et al*., 2017) time records and by a Python function, and with or without the additional time required for post-processing the output files (i.e. parsing relevant information), with varying number of reads (Fig.S2c). The computations were performed using a computing cluster; a single core with 2GB memory was used both for HMC sampling and ADVI approximation (although HMC sampling could be more efficiently run with one core for each MCMC chain hence run time was based on the slowest chain).

### 4 Comparing accuracy in differential methylation analysis

Accuracy in determining differential methylation was measured by generating data sets consisting of two groups (A and B) with varying Δ*θ* (*θ_A_* and *θ_B_* levels are shown in top panels of Fig. S3e) and when one or two of three replicates have low BCR (‘GGB’ and ‘GBB’, respectively). Each group consisted of four biological replicates wherein each biological replicate had three technical replicates each (with different sequencing read coverage, *N*_BS_ = 10 or *N*_BS_ = 6) described in the main text and in Fig. S3a (where *θ*~ Beta(*α,β*), with distribution parameters set to values shown in Fig. S2a).

**Figure S3:**
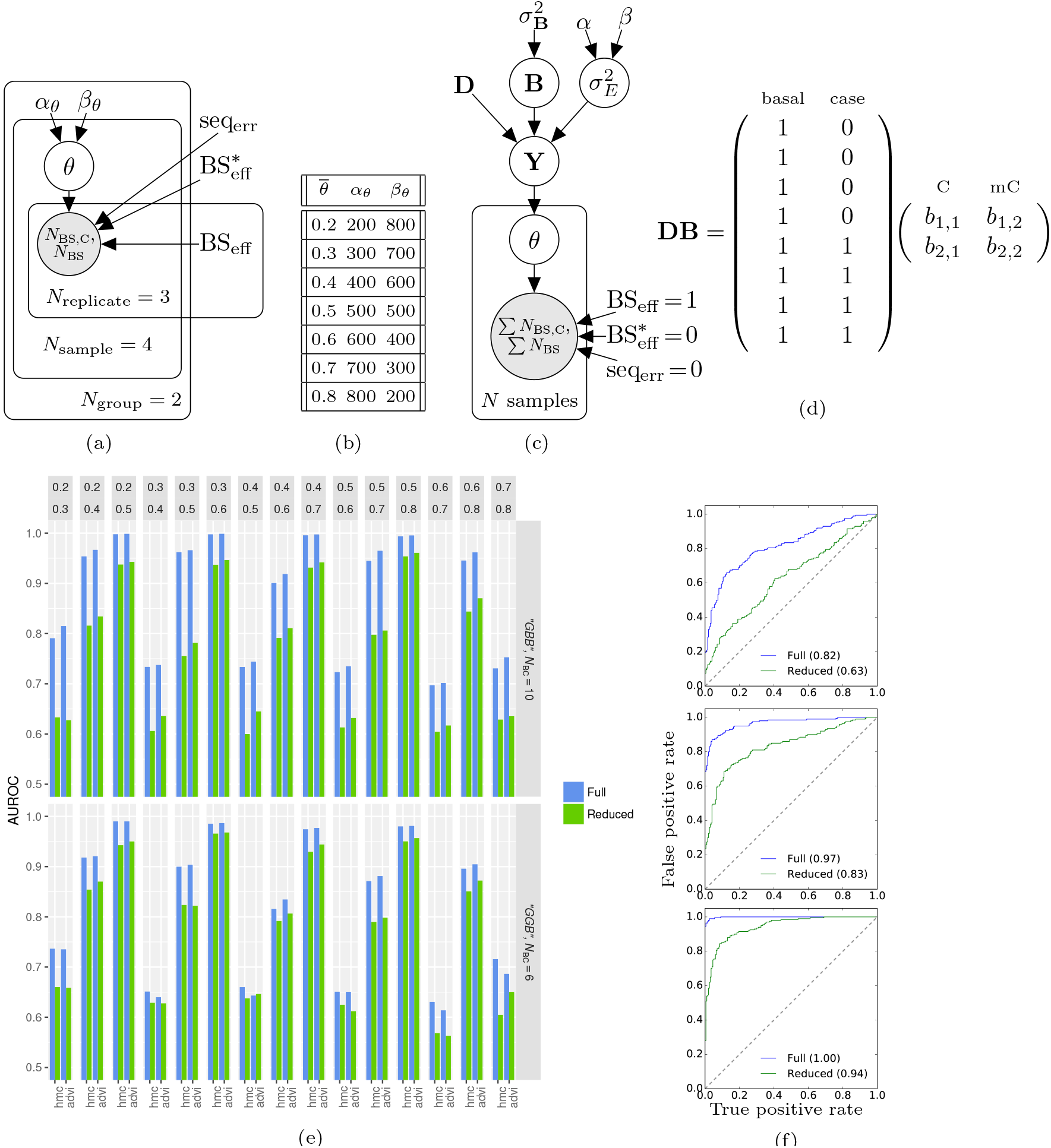
Plate diagram of model for generating dummy data for differential methylation analysis (a), beta parameters used in generating *θ* (b), plate diagram of the reduced LuxRep model (c), design matrix and parameter matrix (d), AUROCs of differential methylation calls (e), and select ROC curves (f). In (c), Σ*N*_BS,C_ and Σ*N*_BS_ denote sums over technical replicates of “C”s and total read counts, respectively.

Differential methylation was analysed using the full and reduced LuxRep models (see Figs. 1a and S3c, respectively, and, for additional details of hyper-priors used,Äijö *et al*. (2016)) evaluated with HMC and ADVI (x-axis in Fig. S3a). Fig. S3d shows the design matrix **D** and parameter matrix **B** used in the general linear model component (Bayes factors were computed using the Savage-Dickey density ratio estimator using samples of *b*_2,1_ and *b*_2,2_, *S* = 1600 and *S* = 1000 from the posterior distributions approximated with HMC and ADVI, respectively). AUROCs were calculated based on ~ 200 positive (Δ*θ*≠0) and ~ 200 negative (Δ*θ* = 0) samples. Select ROC curves generated from the full and reduced models (with technical replicates ‘GBB’ and ‘G’, respectively, and posterior approximation with variational inference) are shown in Fig. S3f where *θ*_1_ = 0.2 and *θ*_2_ was set to 0.3, 0.4 and 0.5 (top, middle and bottom panels, respectively).

### 5 Choosing parameters for variational inference

There are a few parameters which can be tuned to make the ADVI algorithm (Kucukelbir *et al*., 2015) fast but accurate. These parameters are the number of samples used in Monte Carlo integration approximation of expectation lower bound (ELBO), the number of samples used in Monte Carlo integration approximation of the gradients of the ELBO, and the number of samples taken from the approximative posterior distribution. The default values for gradient samples *N_G_* and ELBO samples *N_E_* are 100 and 1 respectively. Here we compare the computation times and accuracy of the differential expression analysis computed using HMC and ADVI with different *N_E_* and *N_G_* values. The tested values for *N_E_* were 100, 200, 500 and 1000 and for *N_G_* 1, 10 and 100. To make the HMC and ADVI methods comparable, the total number of samples retrieved from the approximative posterior distribution is set to 6000 for both methods. To choose the best number of gradient samples and ELBO samples, simulation tests on LuxGLM model were executed. These tests were conducted in the following way: First, simulated data from the LuxGLM model was generated. The number of reads and replicates were varied (the tested values were 6, 12, 24 reads and 6, 10, 20 biological replicates, respectively) and for each combination data sets were generated. The calculation of the Bayes factors was made using different *N_E_* and *N_G_* values. For each setting 200 data sets with differential methylation and without differential methylation were simulated and Bayes factors were calculated. Using the computed Bayes factors, ROC curves and AUROC statistics were produced. Also, the computation times for each parameter value combination were taken down. The results of these tests for the case of 12 reads and 10 replicates are shown in figures S4 and S5.

**Figure S4:**
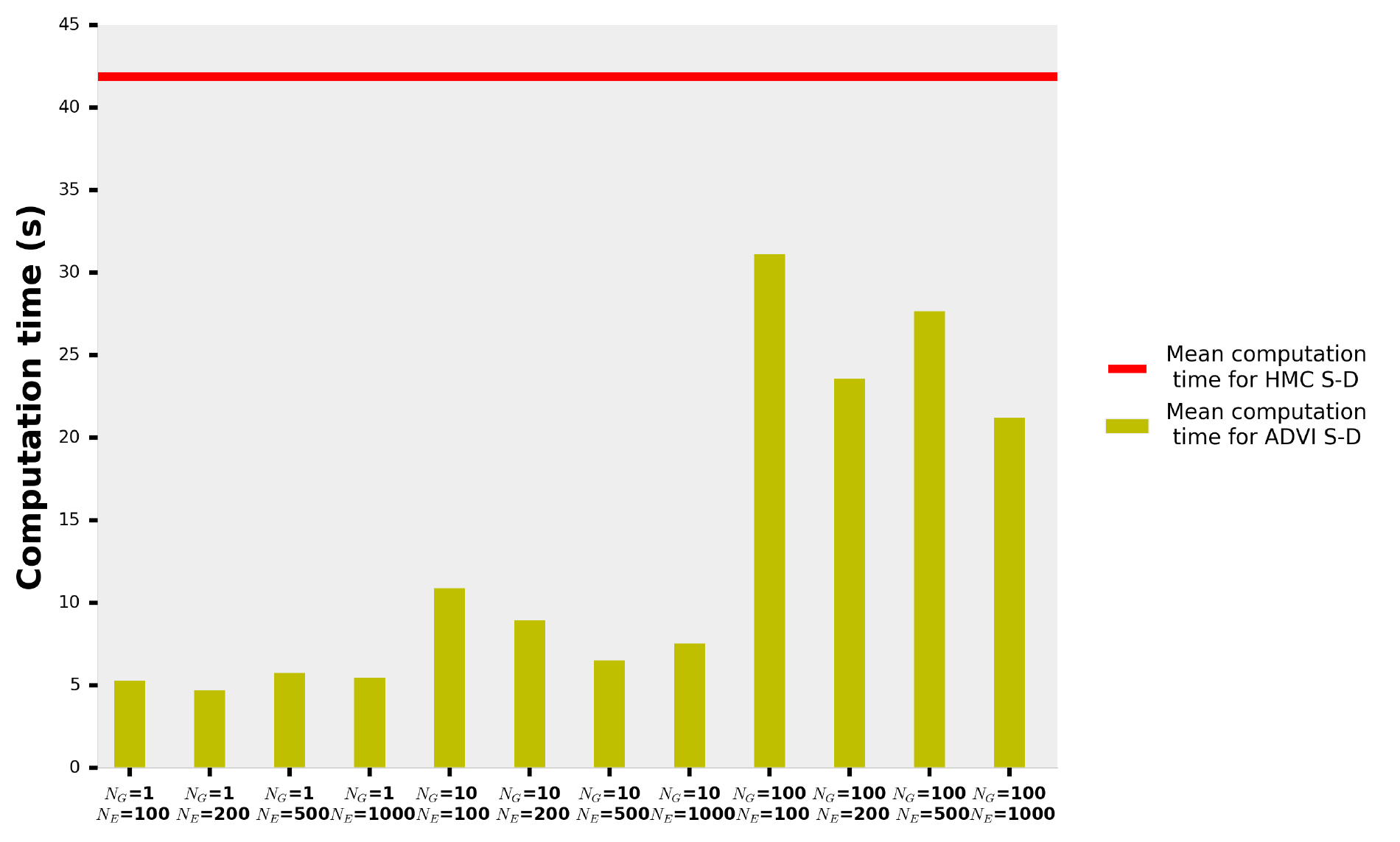
A barplot of the computation times for ADVI with different parameters *N_E_* and *N_G_*. The red line shows the mean computation time with HMC sampler.

**Figure S5:**
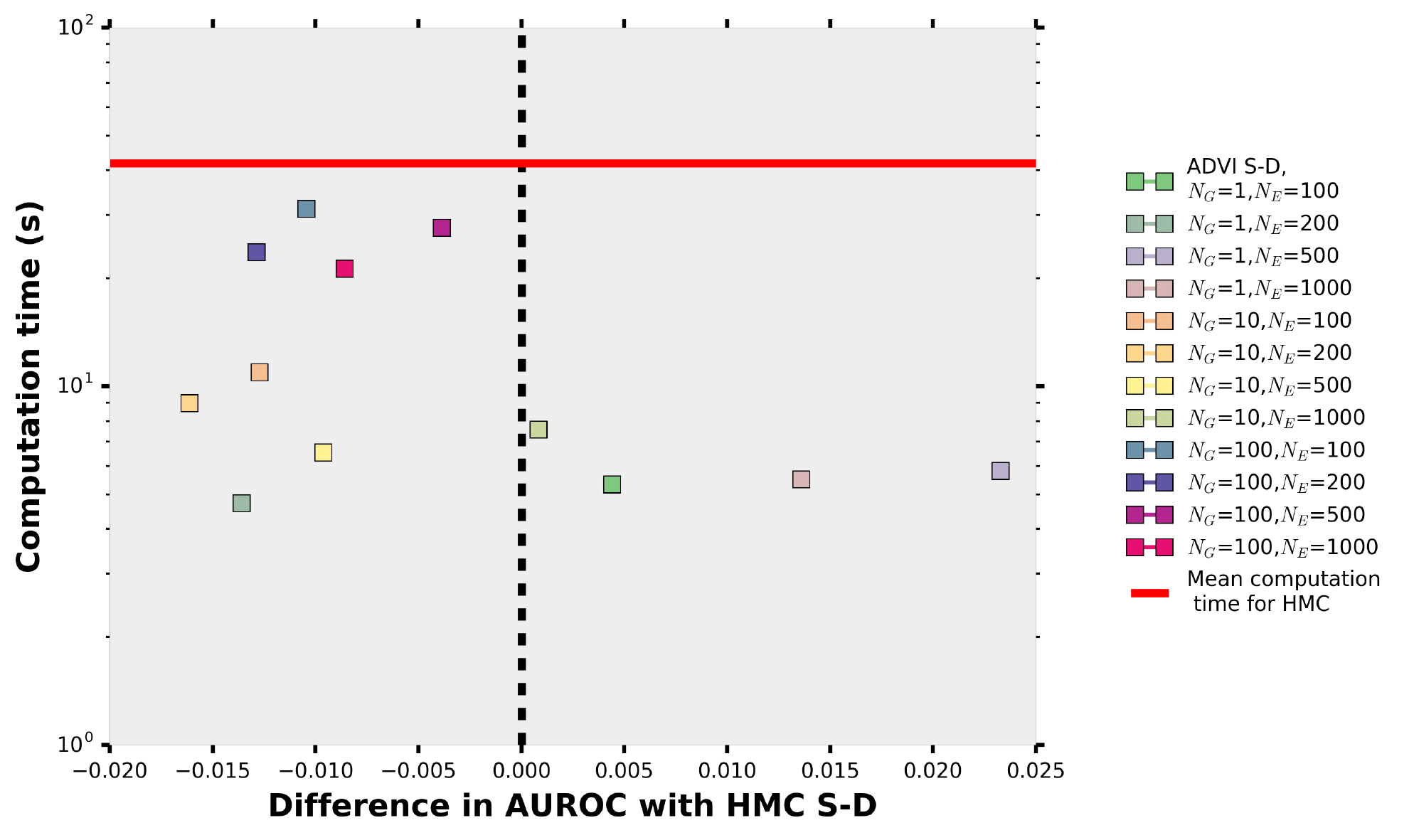
A scatterplot of the computation times with ADVI as function of difference of AUROC values. The different parameter parameter values *N_E_* and *N_G_*. The red line shows the mean computation time for HMC sampler. The dots on the left side of the black dotted line show higher accuracy than Savage-Dickey calculated with HMC and the dots on the right side show lower accuracy than HMC Savage-Dickey. The computation times are plotted in logarithmic scale, and it can be seen that the computation times for ADVI are one magnitude smaller than for HMC when using a good choice of parameters. The graph suggests that ADVI is almost as precise or even more precise as HMC even when the computations are done considerably faster than with HMC.

In Fig. S4 the computation times for different parameter values are shown. In Fig. S5 computation time was plotted as a function of accuracy of the method when compared to the HMC approach. The average computation time for the HMC method is plotted in red. From the figures we can see that with all tested parameter combinations computing Savage-Dickey estimate with ADVI is faster than with HMC. In Fig. S5, on the left side of the dashed line are the parameter combinations which gave better precision than HMC approach.

